# Monocytes and macrophages, targets of SARS-CoV-2: the clue for Covid-19 immunoparalysis

**DOI:** 10.1101/2020.09.17.300996

**Authors:** Asma Boumaza, Laetitia Gay, Soraya Mezouar, Aïssatou Bailo Diallo, Moise Michel, Benoit Desnues, Didier Raoult, Bernard La Scola, Philippe Halfon, Joana Vitte, Daniel Olive, Jean-Louis Mege

## Abstract

To date, the Covid-19 pandemic affected more than 18 million individuals and caused more than 690, 000 deaths. Its clinical expression is pleiomorphic and severity is related to age and comorbidities such as diabetes and hypertension. The pathophysiology of the disease relies on aberrant activation of immune system and lymphopenia that has been recognized as a prognosis marker. We wondered if the myeloid compartment was affected in Covid-19 and if monocytes and macrophages could be infected by SARS-CoV-2. We show here that SARS-CoV-2 efficiently infects monocytes and macrophages without any cytopathic effect. Infection was associated with the secretion of immunoregulatory cytokines (IL-6, IL-10, TGF-β) and the induction of a macrophagic specific transcriptional program characterized by the upregulation of M2-type molecules. In addition, we found that *in vitro* macrophage polarization did not account for the permissivity to SARS-CoV-2, since M1-and M2-type macrophages were similarly infected. Finally, in a cohort of 76 Covid-19 patients ranging from mild to severe clinical expression, all circulating monocyte subsets were decreased, likely related to massive emigration into tissues. Monocytes from Covid-19 patients exhibited decreased expression of HLA-DR and increased expression of CD163, irrespective of the clinical status. Hence, SARS-CoV-2 drives circulating monocytes and macrophages inducing immunoparalysis of the host for the benefit of Covid-19 disease progression.

## Introduction

The novel severe acute respiratory syndrome coronavirus 2 (SARS-CoV-2/2019-n-CoV) emerged in Wuhan (China) at the end of 2019 and caused the coronavirus disease of 2019 (Covid-19) pandemic in a few weeks, affecting more than 18 million people and killing more than 690,000 to date (1). The disease is characterized by a strikingly heterogeneous clinical presentation and prognosis. Most patients are pauci-symptomatic or have fever, cough and fatigue, while a minority experience progression to an acute respiratory distress syndrome or other critically severe conditions. The severity of the disease is related to underlying conditions such as hypertension, diabetes, coronary heart diseases or obesity (2). The mechanisms of the disease remain elusive at this stage, but evidence for a prominent role of the immune system is accumulating. The severity of Covid-19 pneumonia is associated with lymphopenia and a cytokine release syndrome (CRS) (3), which contributes to the massive migration of T cells into tissues, mainly the lung as revealed by accumulation of T cells within lesions (4).

There is evidence that myeloid cells may be involved in the pathophysiology of coronavirus infection, either directly, as a targets for the virus, or indirectly, as effectors of the CRS (5). Indeed, it is known from previous coronavirus outbreaks that macrophages are susceptible to MERS-CoV and SARS-CoV-1 infection (6). Recently, macrophage and monocyte accumulation in the alveolar lumen has been shown in a humanized mice model of SARS-CoV-2 expressing human angiotensin-converting enzyme 2 (ACE2) (7). In addition, SARS-CoV-2 nucleocapsid protein has been detected in lymph nodes and spleen-associated CD169^+^ macrophages from Covid-19 patients (8). Finally, single cell RNA sequencing of pulmonary tissue from Covid-19 patients revealed an expansion of interstitial macrophages and monocyte-derived macrophages (MDM) but not of alveolar macrophages (9). However, whether circulating monocytes and/or macrophages are targets of SARS-CoV-2 and whether monocyte diversity is altered in Covid-19 patients require specific investigation since most studies are based on this hypothesis.

Monocytes are innate hematopoietic cells that maintain vascular homeostasis and ensure early responses to pathogens during acute infections. Three distinct human monocyte subsets are described, based on the expression of CD14 and CD16 surface antigens: classical CD14^+^CD16^-^ monocytes, intermediate CD14^+^CD16^+^ monocytes, and non-classical CD14^-^ CD16^+^ monocytes (0,11). Recently, it has been shown in murine models that classical monocytes are the precursors of non-classical monocytes (12). There is evidence that monocyte subsets exhibit a certain degree of functional specialization. During bacterial infection, classical monocytes are recruited to the sites of inflammation, where they exert typical phagocytic functions and can differentiate into inflammatory dendritic cells or macrophages. Non-classical monocytes crawl along vasculature and surveil the vascular tissue (13). Alterations of monocyte subset frequency have been reported in infectious and inflammatory diseases (10). While macrophages largely arise from monocytes in acute situations such as infection, under homeostatic conditions most tissue macrophages are of embryonic origin and monocytes merely renew this population (14). Consequently, the mobilization of immune cells in Covid-19 might lead to macrophage populations of multiple origin in tissue lesions.

We show here that SARS-CoV-2 has the ability to infect human monocytes and macrophages. SARS-CoV-2 infection stimulated the production of immunoregulatory cytokines, interleukin (IL)-6 and IL-10 in both cell types and triggered in macrophages an original transcriptional program enriched with M2-type genes. Macrophage polarization did not account for permissivity to the virus since M1-and M2-polarized cells were similarly infected by SARS-CoV-2. In Covid-19 patients, the numbers of classical, intermediate and non-classical monocytes were decreased, irrespective of the level of severity. Their expression of CD163, a molecule associated with the immunoregulatory phenotype, was significantly higher than in healthy controls, whereas that of HLA-DR was decreased. Hence, SARS-CoV-2 drives circulating monocytes and macrophages, inducing immunoparalysis of the host for the benefit of Covid-19 disease progression.

## Results

### SARS-CoV-2 infects monocytes and macrophages and stimulates cytokine release

It has been shown that monocytes and macrophages express receptors for SARS-CoV-2 (24), suggesting that the virus targets myeloid cells. We wondered whether SARS-CoV-2 was able to infect human monocytes and macrophages. Monocytes, MDM and Vero cells were incubated with SARS-CoV-2 strain IHU-MI3 (0.1 MOI) for 24 and 48 hours and infection level was measured by RT-PCR and immunofluorescence. SARS-CoV-2 infected efficiently Vero cells (Ct=18.69) after 24 hours, but a lytic process prevented the measurement of viral replication, (**Figure 1A**). Monocytes were also infected after 24 hours (Ct=22.44), but the viral load remained constant thereafter (Ct=22.2) (**Figure 1A**). Similarly, macrophages were efficiently infected with the SARS-CoV-2 strain IHU-MI3 after 24 (Ct=22.49) and 48 hours (Ct=19.67). In contrast to Vero cells, monocytes and macrophages were not uniformly infected, as observed by confocal microscopy (**Figure 1A, right panel**). We next addressed the ability of the IHU-MI3 strain of SARS-CoV-2 to induce the release of soluble mediators from monocytes and MDMs. IL-1β, IL-6, IL-10, TNF-α, IFN-β and TGF-β1 levels were measured in supernatants of monocytes or MDM stimulated with SARS-CoV-2 for 24 and 48 hours. IL-6, IL-10, and IL-1β levels were significantly increased in stimulated monocyte supernatants as compared to unstimulated conditions after 24 hours (**Figure 1B**) and were persistently increased after 48 hours (**Figure 1C**), whereas no difference was observed for TNF-α. In MDM supernatants, levels of IL-6 and IL-10 were increased after 24 and 48 hours (**Figure 1B and C**). TGF-β levels were significantly increased in supernatants from monocytes and MDMs after 48 hours of stimulation. IFN-β was never detected in supernatants from monocytes or macrophages stimulated by SARS-CoV-2 (**Figure 1B and C**). Taken together, SARS-CoV-2 infects monocytes and macrophages. The virus stimulates the release of both pro-and anti-inflammatory cytokines.

**Figure 1.**
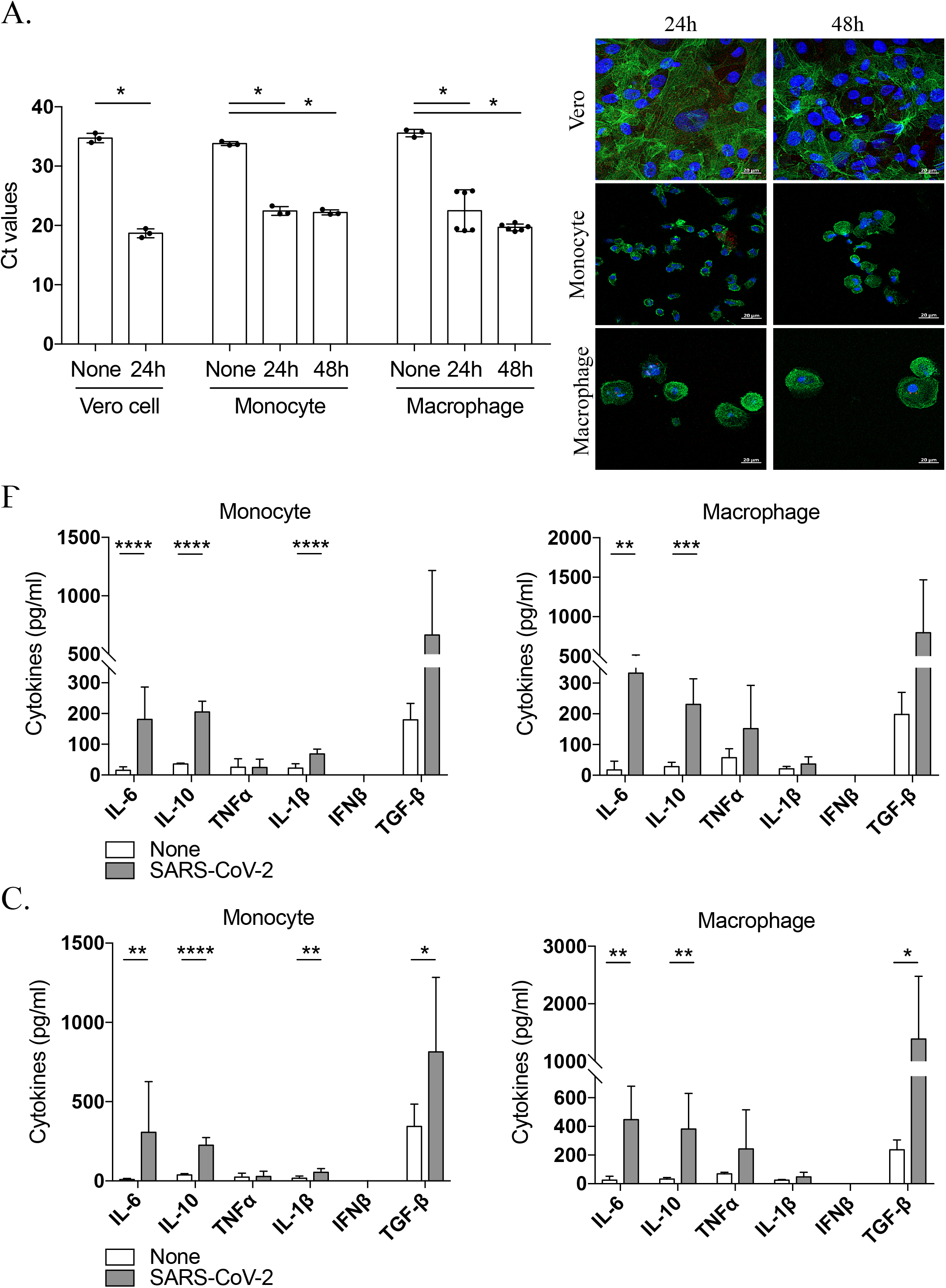
SARS-CoV-2 infects monocytes and macrophages and stimulates cytokine release. Vero E6 cells, monocytes and monocyte-derived macrophages were infected with SARS-CoV-2 IHU-MI3 strain (0.1 MOI) for 24 or 48 hours. (**A**) SARS-CoV-2 quantification was evaluated by RT-PCR, expressed as Ct values and observed in red in infected cells, with the nucleus in blue and F-actin in green (n = 3). Pictures were acquired using a confocal microscope (63x). \*\*\*\**P* < 0.0001 using two-way ANOVA and Turkey’s test for post-hoc comparisons. (**B, C**) Pro-(IFN-β, IL-6, TNF-α, IL-1β) and anti-inflammatory (TGF-β, IL-10) cytokines release was evaluated for SARS-CoV-2-infected monocytes and macrophages at (**B**) 24 and (**C**) 48 hours (n=6). Values represent mean ± standard error of the mean. \**P* < 0.05, \*\**P* < 0.01, \*\*\**P* < 0.001 and \*\*\*\**P* < 0.0001 using Mann-Whitney *U* test.

### SARS-CoV-2 elicits a specific transcriptional program in macrophages

Next, the expression of genes involved in the inflammatory response (*IFNA, IFNB, IFNG, TNF, IL1B, IL6, IL8, CXCL10*) or immunoregulation, (*IL10, TGFB1, CD163*) was measured by qRT-PCR in monocytes and MDM incubated with the virus for 24 and 48 hours. PCA of gene expression using ClustVis software showed that unstimulated and SARS-CoV-2-stimulated monocytes exhibited superimposable programs; in contrast, unstimulated and SARS-CoV-2-stimulated transcriptional programs were clearly distinct in macrophages (**Figure 2A**). In monocytes, a 24 hour-incubation with SARS-CoV-2 stimulated the expression of the whole gene panel, but only *IFNA* gene variation reached the significance level (**Figure 2B**). After 48 hours, expression of *IFNA* declined to levels of unstimulated cells and there was no other change in gene expression (**Suppl figure 1**), suggesting that SARS-CoV-2 was only able to activate gene expression in monocytes in a transient manner.

**Figure 2.**
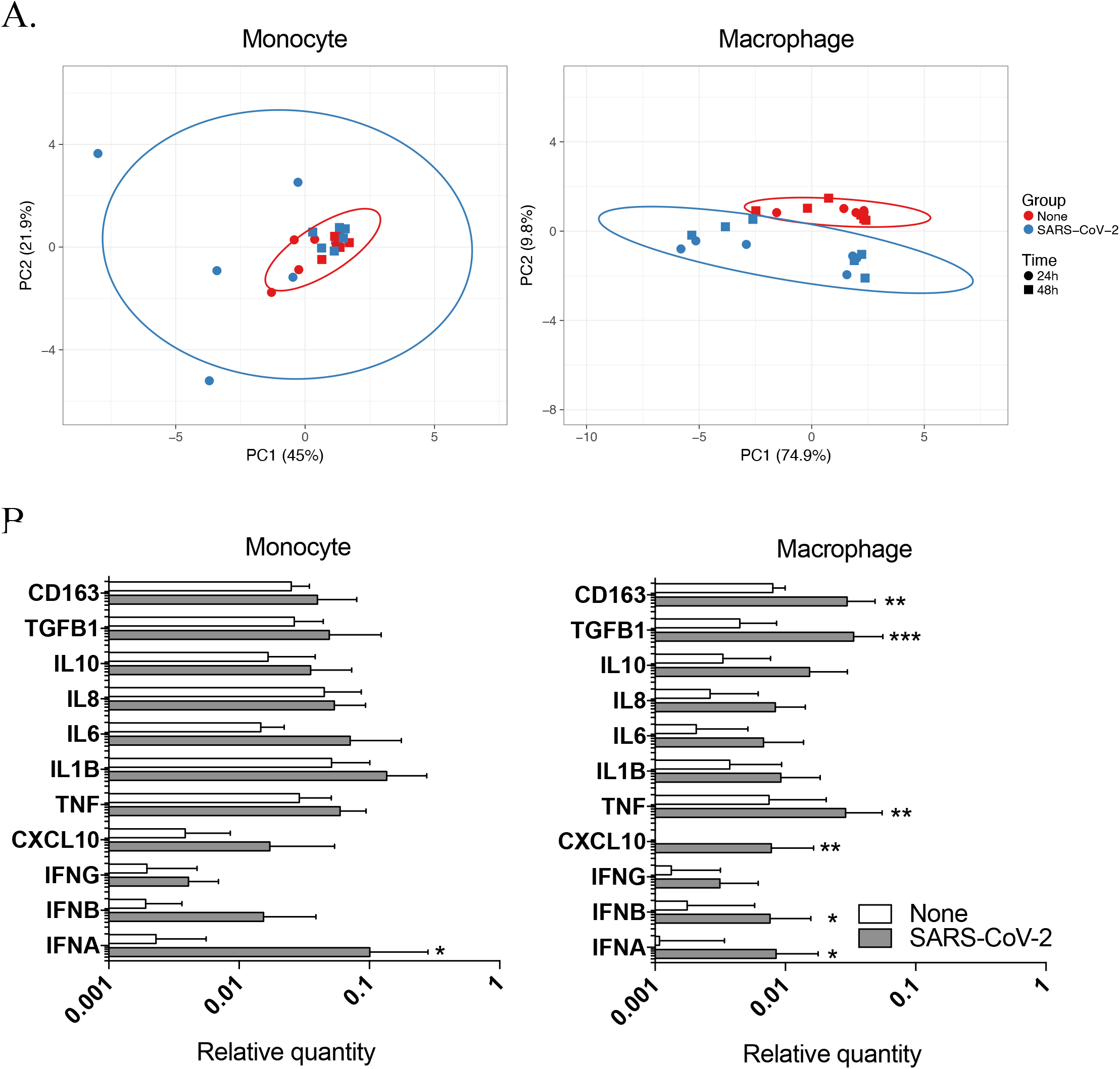
SARS-CoV-2 elicits a specific transcriptional program in macrophages. Monocytes and macrophages were stimulated with SARS-CoV-2 IHU-MI3 strain (0.1 MOI) for 24 or 48 hours (n = 6). The expression of genes involved in the inflammatory response (*IFNA, IFNB, IFNG, TNF, IL1B, IL6, IL8, CXCL10*) or immunoregulation (*IL10, TGFB1, CD163*) was investigated by qRT-PCR after normalization with housekeeping actin gene as endogenous control. (**A**) Data are illustrated as principal component analysis obtained using ClustVis webtool for uninfected and SARS-CoV-2 infected cells in red and blue respectively, with round points for 24 hours and square points for 48 hours of stimulation. (**B**) Relative quantity of investigated genes at 24 hours of stimulation was evaluated for monocytes (left panel) and macrophages (right panel). Values represent mean ± standard error of the mean. \**P* < 0.05, \*\**P* < 0.01 and \*\*\**P* < 0.001 using two-way ANOVA and Sidak’s test for post-hoc comparisons.

We next investigated the transcriptional program induced by SARS-CoV-2 in macrophages. Similar to monocytes, at 24 hours, SARS-CoV-2 significantly increased the expression of antiviral (*IFNA* and *IFNB*), inflammatory (*CXCL10* and *TNF*), and immunoregulatory genes (*TGFB1* and *CD163*) (**Figure 2B**). The increase in gene expression was no longer observed after 48 hours except for *TGFB* and *CD163* (**Suppl figure 1**). Hence, the early transcriptional program of infected macrophages consisted of genes associated with M1 profile (type I *IFN, CXCL10*) and M2 profile (*TGFB1* and *CD163*), suggesting that SARS-CoV-2 does not induce clear polarization of macrophages at the onset of the infection but rather a delayed shift toward a M2-type.

### Macrophage polarization and SARS-CoV-2 infection

As SARS-CoV-2 induced an early M1/M2 followed by a late M2 program in macrophages, we investigated the effect of macrophage polarization status on infection. MDM polarization was induced by IFN-γ (20 ng/ml) and lipopolysaccharide (100 ng/ml) (M1), IL-4 (20 ng/ml) (M2), or was kept at a resting state without polarization (M0). The polarization status was confirmed by measuring the expression of M1 (*IL1B, IL1RA, IL6, IL12, CXCL10, TNF, NOS2, IFNG*) and M2 genes (*ARG1, IL10, MR, CD163, TGFB*). PCA and hierarchical clustering confirmed the induction of three distinct activation statuses (**Suppl. figure 2**). The expression of polarization-related genes was investigated after 24 and 48 hours of SARS-CoV-2 stimulation of M1 and M2 polarized macrophages. The hierarchical clustering showed that unstimulated and SARS-CoV-2-stimulated MDM (M0, M1 and M2) were present on two distinct branches but the discrimination of responses as a function of polarization was not possible (**Suppl figure 3**). Regarding pro-(IL-6, TNF) and anti-(IL-10, TGF-β) inflammatory cytokines, SARS-CoV-2 stimulation significantly increased the release of both cytokine groups in M1-and M2-polarized macrophages after 24 and 48 hours (**Figure 3A, B**). In addition, no differences were observed in the viral load of M0, M1 or M2 macrophages (**Figure 3C**). We next performed the same experiment using M0, M1 or M2 THP-1 macrophages. The choice of a cell line instead of primary macrophages aimed at minimizing inter-individual variations. When THP-1 macrophages were M1 polarized, the viral load was similar to that of non-polarized (M0) macrophages. In contrast, SARS-CoV-2 load was significantly decreased in M2 polarized macrophages as compared with M0 macrophages (**Figure 3C**). Although a type 2 immune response was associated with lesser infection of macrophages, their polarization did not appear critical for SARS-CoV-2 infection.

**Figure 3.**
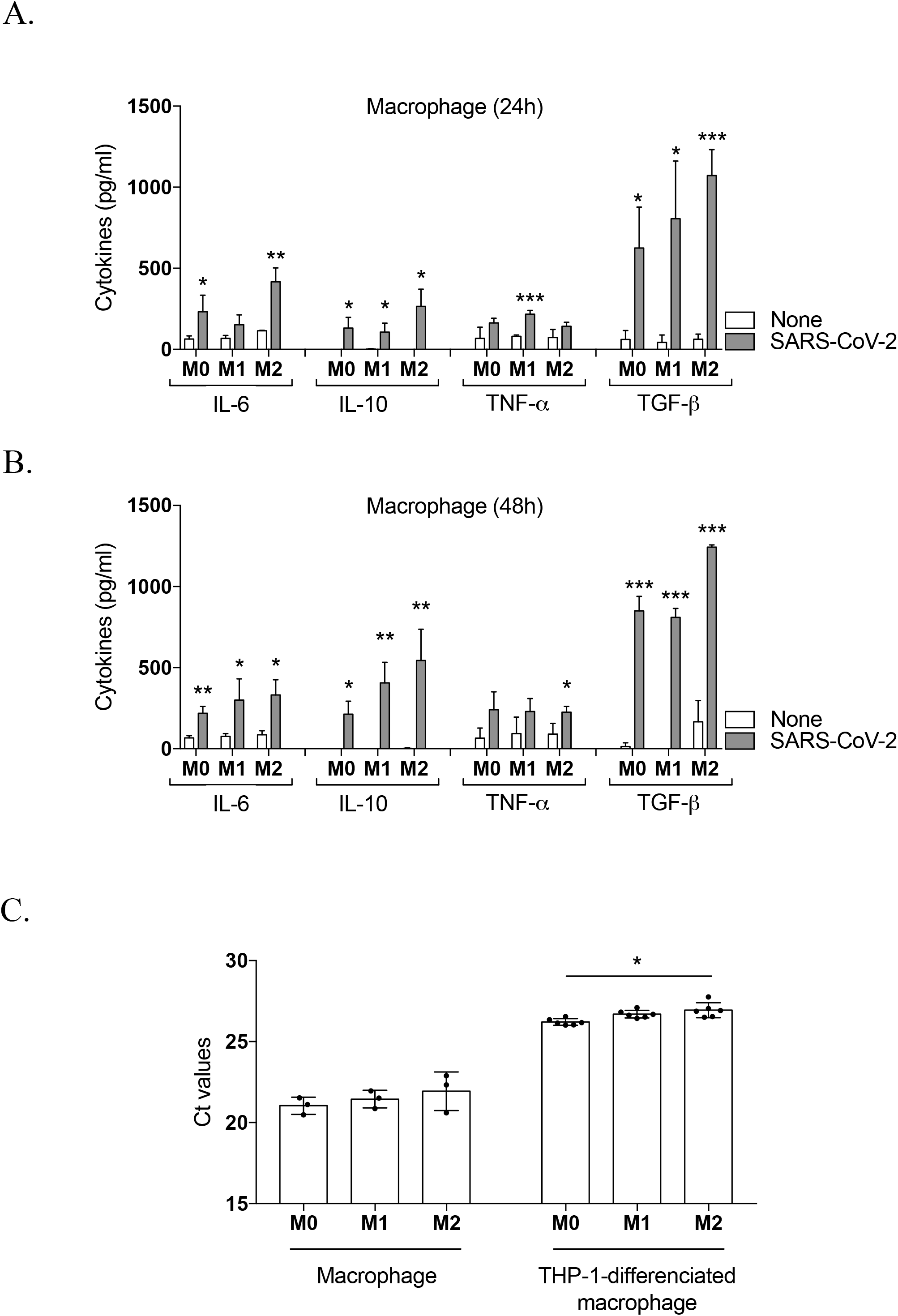
Investigation of polarized macrophages in the SARS-CoV-2 response. Macrophages and PMA-differentiated THP-1 cells were polarized by treatment with IFN-γ (20 ng/ml) and lipopolysaccharide (100 ng/ml) (M1), IL-4 (20 ng/ml) (M2) or without agonist (M0). Polarized macrophages were stimulated for (**A**) 24 or (**B**) 48 hours with IHU-MI3 SARS-CoV-2 strain and IL-6, IL-10, TNF-α and TGF-β release were evaluated in the culture supernatants by ELISA (n = 3). \**P* < 0.05, \*\**P* < 0.01 and \*\*\**P* < 0.001 using Mann-Whitney *U* test. (**C**) Virus quantification was assessed by the evaluation of the Ct values for polarized SARS-CoV-2-infected macrophages (n = 3) and PMA-differentiated THP-1 cells (n = 6) at 24 hours post-infection. Values represent mean ± standard error of the mean. \**P* < 0.05 using two-way ANOVA and Turkey’s test for post-hoc comparisons.

### Monocyte subsets are altered in SARS-CoV-2-infected patients

Following the demonstration of a direct *in vitro* effect of SARS-CoV-2 on monocytes and macrophages, we wondered if the frequency of monocyte subsets was affected in Covid-19 patients. Monocyte subsets were analyzed for CD14, CD16 and HLA-DR expression by flow cytometry in 76 Covid-19 patients and compared to healthy blood donors (**Figure 4A**). In the latter, classical monocytes were the best represented monocyte subset (9.17% of total PBMCs), while intermediate and non-classical monocytes accounted for 0.42% and 0.60%, respectively. In Covid-19 patients, the percentages of classical (2.03%), intermediate (0.23%) and non-classical monocytes (0.22%) were significantly lower than in healthy controls (**Figure 4A**). Hence, the monocytopenia previously reported in patients infected with SARS-CoV-2 (25) affected all three monocyte subsets. We wondered if circulating monocytes displayed changes in the expression level of activation-associated membrane markers. Hence, we measured the expression of HLA-DR, a canonical marker of monocyte activation, and CD163, an immunoregulatory marker. As shown in **figure 4B**, all three monocyte subsets expressed HLA-DR and CD163. In Covid-19 patients, the expression level of HLA-DR was significantly decreased in intermediate and non-classical monocytes, whereas it remained similar to controls in classical monocytes (**Figure 4B**). In contrast, the expression of CD163 was significantly increased in classical and non-classical monocytes (**Figure 4B**). The opposite effect of Covid-19 on HLA-DR and CD163 expression suggests that their activation status was shifted to an immunoregulatory program. This phenotypic profile of patient monocytes was partly recapitulated by incubating control monocytes with SARS-CoV-2. SARS-CoV-2 increased HLA-DR expression in a dose dependent manner after 24 hours and decreased it after 48 hours. The inverted pattern was observed with CD163 (**Figure 4C**). Finally, we wondered if the decrease in monocyte subsets and altered expression of HLA-DR and CD163 reflected the severity of Covid-19. There were no significant differences in monocyte phenotype among mild, moderate, and severe patients (**Figure 5**). Hence variation of monocyte HLA-DR and CD163 expression in Covid-19 patients was induced by SARS-CoV-2 infection, but was not related to subsequent disease severity.

**Figure 4.**
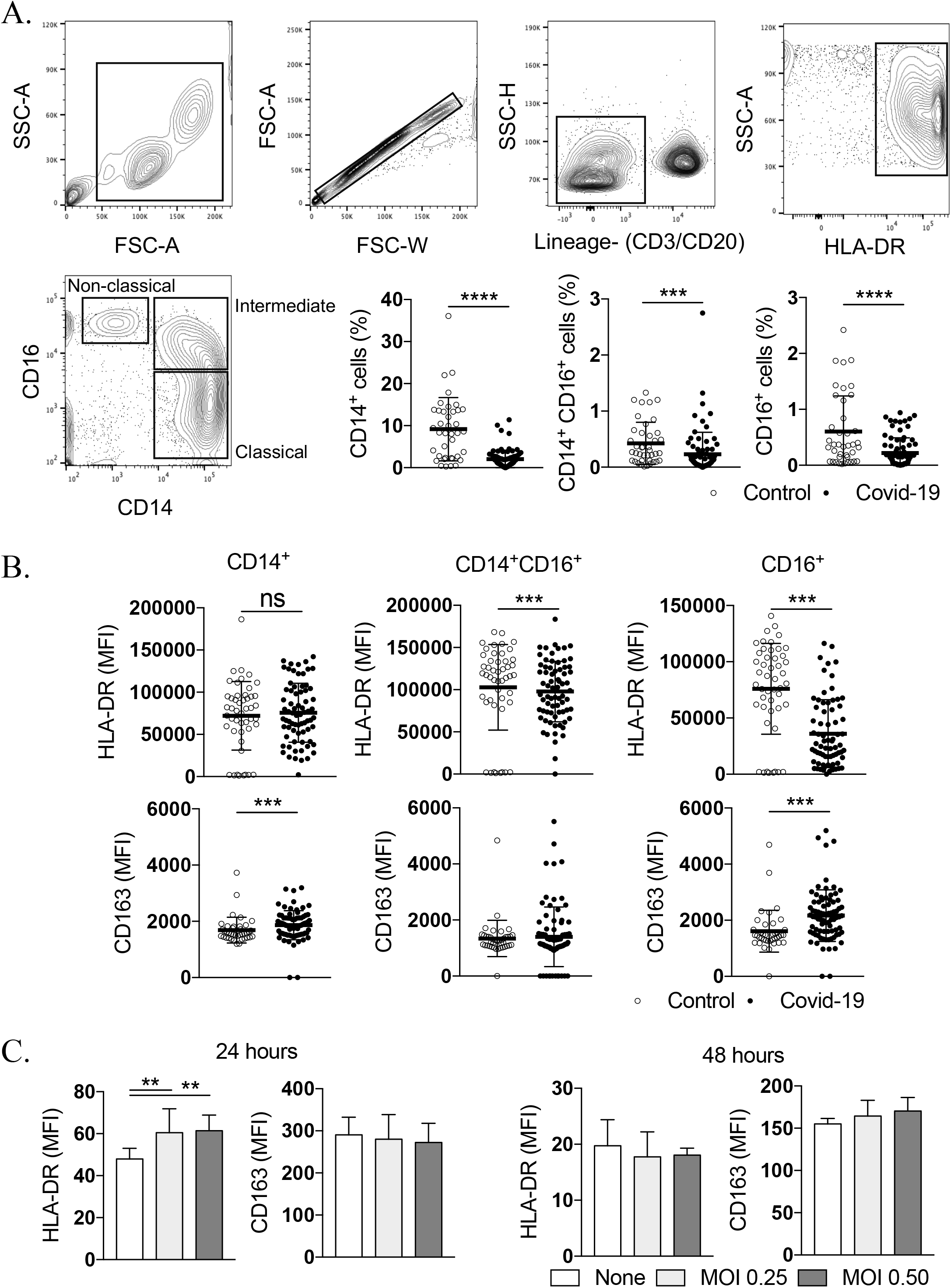
Monocyte subsets are altered in SARS-CoV-2-infected patients. PBMCs from healthy donors and Covid-19 patients were isolated and monocyte sub-populations were investigated by flow cytometry (**A**) Representative flow cytometry plot showing the gating strategy to investigate non-classical, classical and intermediate HLA-DR^+^ monocytes from Covid-19 patients and healthy donors as control. (**B**) Mean fluorescence intensity (MFI) of HLA-DR and CD163 expression was investigated for CD14^+^, CD14^+^/CD16^+^ and CD16^+^ monocyte populations from healthy and Covid-19 patients. (**C**) Monocytes from healthy donors were stimulated with SARS-CoV-2 IHU-MI3 strain (0.25 or 0.5 MOI). The expression of HLA-DR and CD163 was observed at 24 and 48 hours of infection. \*\**P* < 0.01, \*\*\**P* < 0.001 and \*\*\*\**P* < 0.0001 using t-test.

**Figure 5.**
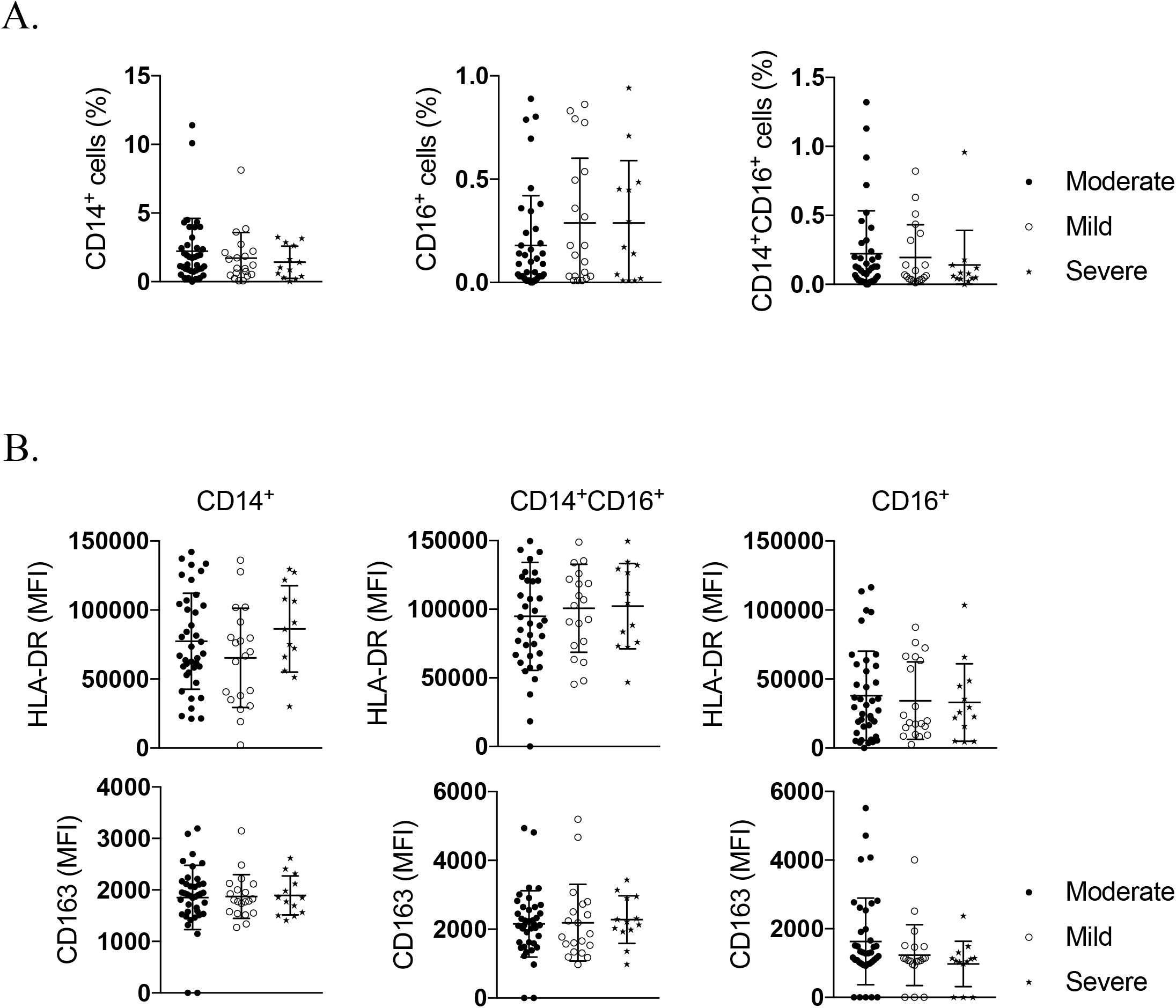
Peripheral blood mononuclear cells from Covid-19 patients were isolated and monocyte sub-populations were investigated by flow cytometry. (**A**) Non-classical, classical and intermediate HLA-DR^+^ monocytes were evaluated from moderate, mild and severe Covid-19 clinical population. (**B**) Mean fluorescence intensity (MFI) of HLA-DR and CD163 expression was investigated for CD14^+^, CD14^+^/CD16^+^ and CD16^+^ monocyte populations from moderate, mild and severe Covid-19 patients using t-test.

## Discussion

We showed that SARS-CoV-2 efficiently infects human monocytes and macrophages. This is reminiscent of previous reports about SARS-CoV-1 that infects human macrophages but does not replicate within (26). In addition, macrophages infected by SARS-CoV-1 were detected in lungs of SARS patients (5). Recently, *post-mortem* examination of lymph nodes and spleen revealed the presence of SARS-CoV-2 nucleocapsid protein in macrophages that express CD169, a maker of macrophages from the splenic marginal zone (8). Using an unsupervised computational pipeline that can detect viral RNA in any scRNA-seq data set, an enrichment of SARS-CoV-2 reads in macrophages expressing secreted phosphoprotein 1 was observed (27). Although monocytes and macrophages express the molecular machinery to recognize and internalize SARS-CoV-2 such as ACE2, TMPRSS2 and ADAM17 (24), the ability of the virus to replicate within these cells is not fully understood. Our results favor the hypothesis of an abortive infection similar to SARS-CoV-1 (28) but clearly distinct from MERS-CoV replication in macrophages (29).

The infection of monocytes and macrophages is associated with the production of inflammatory cytokines that contribute to the CRS described in patients and involved in disease pathogenesis. Monocytes and macrophages exhibit a common secretory profile associating the release of IL-6, IL-10 and TGF-β and the absence of IFN-β. The impaired IFN production is consistent with the reported inhibition of type I IFNs by SARS-CoV-1 and the lack of interferon regulatory factor 3 activation in macrophages and myeloid dendritic cells (5). In addition to preventing IFN-α/β responses, SARS-CoV downregulated IFN-related genes in THP-1 cell lines (30). At least three SARS-CoV proteins, namely N protein, OrfB3 and Orf6, are known to antagonize the IFN-β response (31). While the release of IL-6 is consistent with previous reports on the ability of SARS-CoV-1 to stimulate IL-6 secretion in human MDM, the lack of significant changes in TNF release was not expected (26). It has been shown in an *in-situ* study of *post-mortem* samples that SARS-CoV-2 induces IL-6 more efficiently than TNF (8). In our hands, both monocytes and macrophages released IL-10 and TGF-β, suggesting that anti-inflammatory cytokines are also involved in cell responses to infection. The release of TGF-β by monocytes and macrophages may be associated with tissue repair and generation of fibrosis that complicates Covid-19 evolution (32). Taken together, our results suggest that the early response of monocytes and macrophages is inflammatory whereas the delayed response promotes tissue repair. This model is in line with the immune response unfolding in Covid-19 patients, in whom myeloid cells interact with innate and adaptive immune partners able to redirect immune responses towards an inflammatory status.

We found that SARS-CoV-2 differently affected the transcriptional programs of monocytes and macrophages. In monocytes, SARS-CoV-2 elicited a transient program dominated by the upregulation of IFNα gene, while macrophages exhibited a more diversified transcriptional program associating inflammatory and anti-inflammatory genes, which shifted to an anti-inflammatory program of M2 type. Hence, SARS-CoV-2 affected macrophage polarization according to the kinetics of infection. Previous reports on SARS-CoV-1 infection showed a direct effect of virus on macrophage activation. In an African green monkey model, SARS-CoV-1 activated pulmonary macrophages by polarizing them toward a M1 profile associated with decreased viral load but persistence of inflammation (33). In a murine model of SARS-CoV-1 infection, alveolar macrophages were repolarized to limit T cell activation (34). Another study revealed that SARS-CoV-1 induced non protective M2 polarization in lung macrophages from infected mice(35). Whether macrophage polarization affected their capacity to control SARS-CoV-2 replication was not addressed. Using polarized MDM and differentiated THP-1 cells, we found that non polarized and M1 type polarized macrophages were permissive to SARS-CoV-2. This may explain why obesity and diabetes, conditions associated with M1 macrophage polarization, are critical comorbidities in Covid-19 (36). In our hands, M2 type macrophages tended to be less permissive to SARS-CoV-2. As estrogens favor M2 polarization (37), this may explain why women are less affected than men by Covid-19. In addition, patients with allergic asthma seem to be less susceptible to the virus (38). Our results suggest that, instead of inducing a clear polarization, SARS-CoV-2 exacerbates macrophage responses whatever the type of polarization.

The myeloid compartment was analyzed through monocyte frequencies and the expression of membrane markers. Previous reports established that monocytopenia was detected in Covid-19 patients in association with lymphopenia. We showed that this decrease in circulating monocytes affects all monocyte subsets. There is a lack of consensus about the variations of monocyte count in Covid-19, probably because of the diversity of measurement tools and the heterogeneity of patients in terms of evolution. A.J. Wilk *et al* reported depletion of CD16^+^ monocytes including intermediate and non-classical monocytes in a single-cell RNA sequencing study of Covid-19 PBMCs (39). A Cytof study of CD45^+^ mononuclear cells revealed an initial increase in cell count from mild to severe followed by a decline in more severe patients (40). Expansion of IL-6 producing CD14^+^CD16^+^ monocytes was reported in Covid-19 patients hospitalized in intensive care units (ICU) as compared with patients not requiring ICU care (32).

Besides monocyte depletion in patients, remaining monocytes were characterized by a down-modulation of HLA-DR and upregulation of CD163. HLA-DR down-modulation is in agreement with previous studies. A. Gatti *et al* reported downregulation of monocyte HLA-DR in patients with severe SARS-CoV-2 (41). A sc-RNA seq study revealed that genes encoding class II HLA molecules were downmodulated in Covid-19 patients(39). P. Bost reported a disease-severity-associated signature in MDM in which MHC II and type I IFN genes were downmodulated (27). Another study showed that CD14^+^ monocytes maintained the expression of HLA-DR in mild/moderate patients, with down-modulation occurring only in severe forms (42). Previously unreported CD163 upregulation in Covid-19 patients suggests a monocyte polarization toward a M2-type profile. Immunohistochemical staining of SARS pneumonia demonstrated CD163^+^ M2 macrophages *in situ* (43). M2 polarization is the consequence of the release of immunoregulatory cytokines, but also of the interaction with the virus. Indeed, we showed a trend in monocytes infected with SARS-CoV-2 with an increase in CD163 expression paralleling low HLA-DR expression. It is known that IL-6 antagonizes HLA-DR expression and the addition of the specific inhibitor of IL-6 pathway, tocilizumab, partially restores HLA-DR expression of CD14^+^ monocytes from Covid-19 patients (42). A synergism between SARS-CoV-2 and IL-6 is likely necessary to down-modulate the expression of HLA-DR and to disarm microbicidal competence of monocytes and macrophages.

Here, we showed that SARS-CoV-2 infects monocytes and macrophages without cytopathic effect and induces a more sustained activation program in macrophages. Monocyte and macrophage response to SARS-CoV-2 is more complex than expected from the observation of CRS, to which they poorly contribute. The investigation of circulating monocytes suggested that massive migration to tissues had occurred and remaining blood monocytes exhibit a repairing profile. This observation may help understand the risk of post-Covid-19 complication including fibrosis. Indeed, a subset of macrophages with a pro-fibrotic program has been described in patients with Covid-19 (32). The lack of correlation between monocyte count and monocyte functional polarization with severity stages suggest that monocytes are markers of SARS-CoV-2 infection. It is likely that other membrane markers of myeloid cells are modulated according to disease progression and reflect more accurately the inflammatory context associated with the severity. Taken together, our study showed that monocytes and macrophages are targets of SARS-CoV-2, and their manipulation may open the way for therapeutic perspectives.

## Methods

### Patients and ethical statement

Seventy-six consecutive patients with SARS-CoV-2 infection confirmed through reverse transcriptase-polymerase chain reaction (RT-PCR, Life Technologies, Carlsbad, CA, USA) from March 16 through March 27, 2020 at the University Hospitals of Marseille, France, were included. Not later than 48 hours post-diagnosis, patients underwent clinical laboratory tests and blood was drawn through venipuncture into EDTA anticoagulated tubes. Epidemiological, demographic, clinical, laboratory and outcome data were obtained from a retrospective, non-interventional review of the medical charts and laboratory results. Demographic characteristics of the study population are presented in **table 1**. This study was performed on excess EDTA-anticoagulated total blood samples. According to French law, the patients had received information that their excess samples and clinical data might be used for research purposes, and retained the right to oppose (5,16).

**Table 1.**
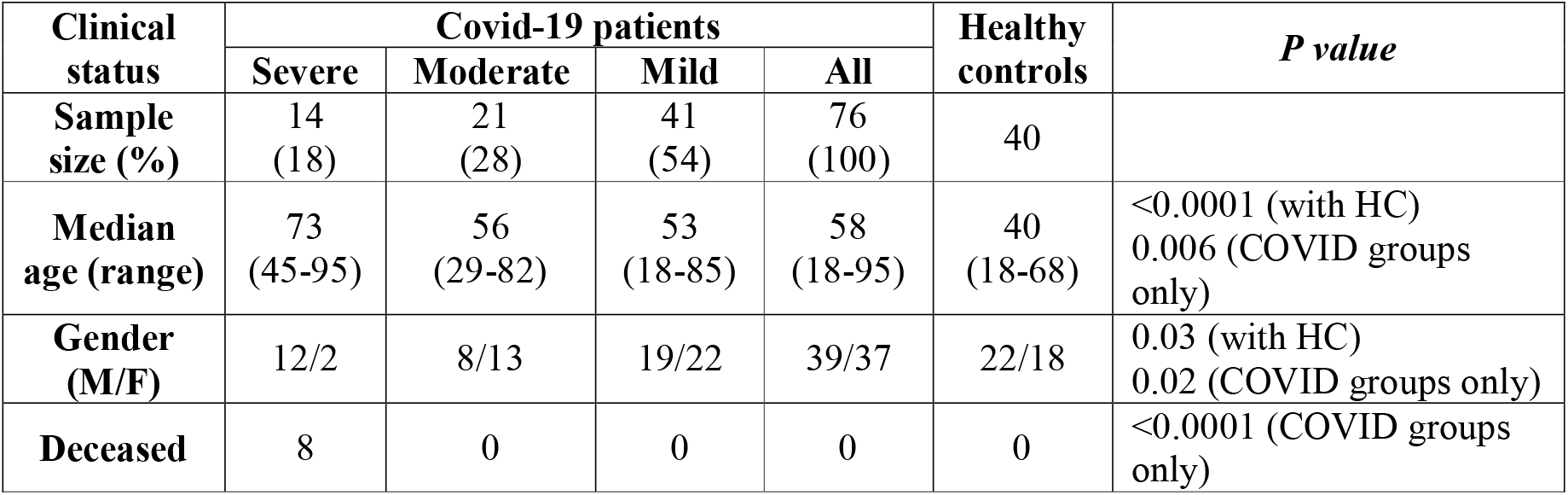
Clinical and demographic data of the study population. Seventy-six consecutive Covid-19 patients and 41 healthy controls were analyzed. Demographic data were available for 40 healthy controls. Nonparametric Kruskall-Wallis test was used for group comparison. HC, healthy control; F, female, M, male.

| Clinical status | Covid-19 patients |  |  |  | Healthy controls | *P value* |
| --- | --- | --- | --- | --- | --- | --- |
|  | Severe | Moderate | Mild | All |  |  |
| Sample size (%) | 14 (18) | 21 (28) | 41 (54) | 76 (100) | 40 |  |
| Median age (range) | 73 (45-95) | 56 (29-82) | 53 (18-85) | 58 (18-95) | 40 (18-68) | <0.0001 (with HC)<br>0.006 (COVID groups only) |
| Gender (M/F) | 12/2 | 8/13 | 19/22 | 39/37 | 22/18 | 0.03 (with HC)<br>0.02 (COVID groups only) |
| Deceased | 8 | 0 | 0 | 0 | 0 | <0.0001 (COVID groups only) |

### Cell isolation

Peripheral blood mononuclear cells (PBMCs) were isolated from the blood of Covid-19 patients and from buffy coats from healthy blood donors (Convention *N°7828*, “Etablissement Français du Sang”, Marseille, France) by density gradient centrifugation using Ficoll (Eurobio, Les Ulis, France) as previously described (17). Monocytes were purified by CD14 selection using MACS magnetic beads (Miltenyi Biotec, Bergisch Glabach, Germany) and cultured in Roswell Park Memorial Institute medium-1640 (RPMI, Life Technologies, Carlsbad, CA, USA) containing 10% inactivated human AB-serum, 2 mM glutamine (Sigma Aldrich, Saint-Quentin-Fallavier, France), 100 U/mL penicillin and 50 µg/mL streptomycin (Life Technologies). After 3 days, the medium was replaced by RPMI-1640 containing 10% fetal bovine serum (FBS, Life Technologies) and 2 mM glutamine, and cells were differentiated into macrophages for 4 additional days. For some experiments, THP-1 macrophages were used and cultured in RPMI-1640 containing 10% FBS, 2mM glutamine and 100 U/mL penicillin and 50 µg/mL streptomycin and differentiated into macrophages after treatment with 50 ng/ml phorbol-12-myristate 13-acetate (PMA, Sigma Aldrich) for 48 hours (8,19).

### Virus production and cell infection

SARS-CoV-2 strain IHU-MI3 was obtained after Vero E6 cells (American type culture collection ATCC® CRL-1586(tm)) infection in Minimum Essential Media (MEM) (Life Technologies) supplemented with 4% FBS as previously described (20).

Cells were infected with 50 µl virus suspension (0.25, 0.5 or 0.1 multiplicity of infection (MOI)) for 24 or 48 hours at 37°C in the presence of 5% CO_2_ and 95% air in a humidified incubator.

### Immunofluorescence

After a 24 or 48-hour infection, cells were incubated in blocking buffer (Phosphate buffer saline (PBS) supplemented with 5% FBS and 0.5% Triton X-100) for 30 minutes and washed before incubation with an anti-SARS-CoV-2 spike protein antibody (Life Technologies). Nuclei and F-actin were stained using DAPI and Phalloidin (Life Technologies) respectively. Pictures were obtained using an LSM800 Airyscan confocal microscope (Zeiss) and a 63X oil objective.

### Viral RNA extraction and q-RTPCR

Viral RNA was extracted from infected cells using NucleoSpin® Viral RNA Isolation kit (Macherey-Nagel, Hoerdt, France) following the manufacturer’s recommendations. Virus detection was performed using One-Step RT-PCR SuperScript(tm) III Platinum(tm) Kit (Life Technologies). Thermal cycling was achieved at 55°C for 10 minutes for reverse transcription, pursued by 95°C for 3 minutes and then 45 cycles at 95°C for 15 seconds and 58°C for 30 seconds using a LightCycler 480 Real-Time PCR system (Roche, Rotkreuz, Switzerland). The primers and the probes were designed against the E gene (20).

### RNA isolation and q-RTPCR

Total RNA was extracted from monocytes or macrophages (2.10^6^ cells/well) using the RNeasy Mini Kit (Qiagen, Courtaboeuf, France) and DNase I treatment to eliminate DNA contaminants (21). The quality and quantity were evaluated using a spectrophotometer (Nanodrop Technologies, Wilmington, USA). Reverse transcription of isolated RNA was performed using a Moloney murine leukemia virus-reverse transcriptase kit (Life Technologies) and oligo(dT) primers. q-PCR was performed using the Smart SYBRGreen fast Master kit (Roche Diagnostics, Meylan, France) and a CFX Touch RTPCR Detection System (Bio-Rad, Marnes-la-Coquette, France) using specific primers (**Table 2)**. The results were normalized using the housekeeping endogenous control *actb* gene encoding β-actin and are expressed as relative expression of investigated genes using the formula 2^-^ΔCt where ΔCt = Ct_Target_ -Ct_Actin_ as previously described (22). The threshold cycle (Ct) was defined as the number of cycles required to detect the fluorescent signal.

**Table 2.**
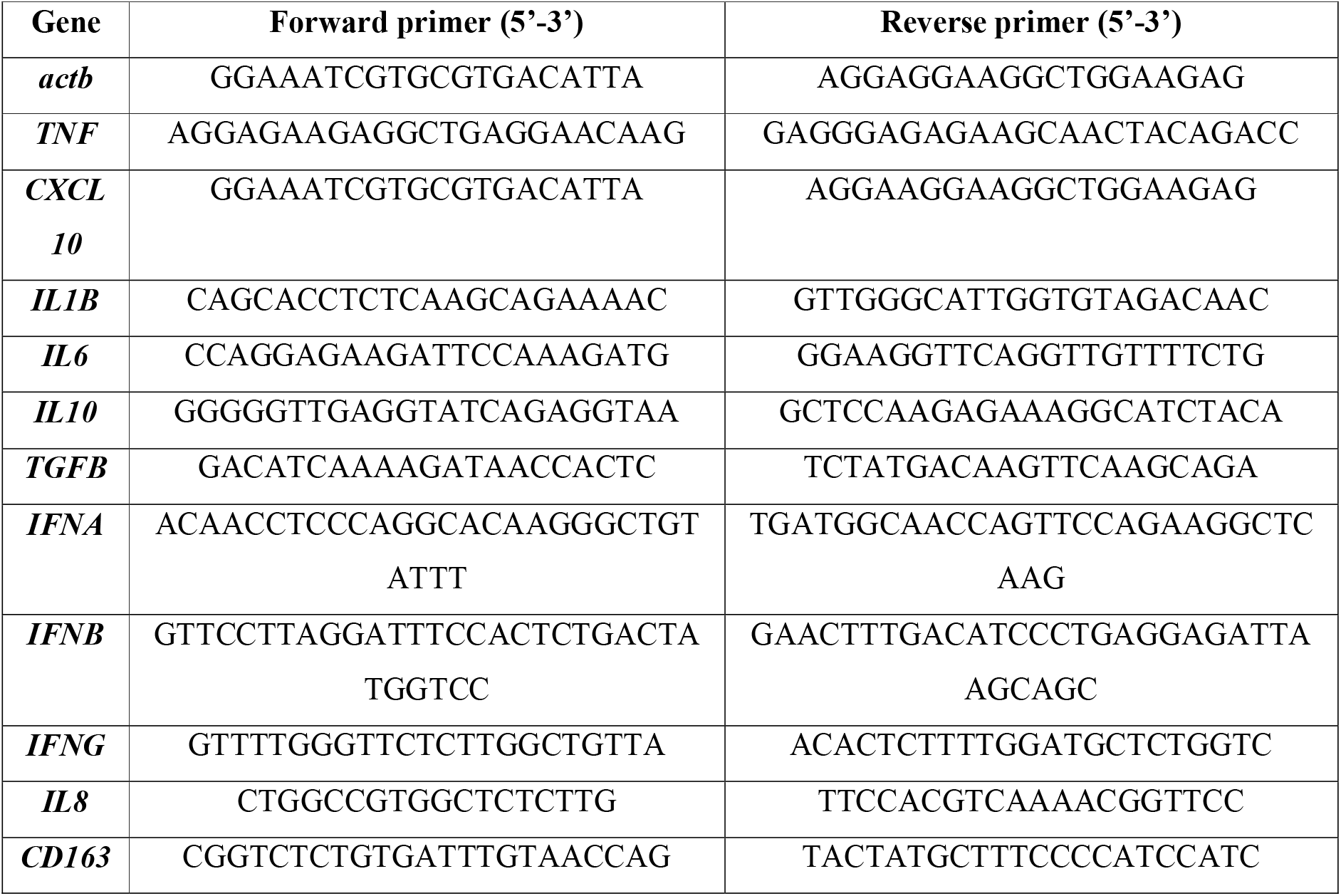
List of primers used for q-RTPCR.

### Immunoassays

Cell supernatants were collected and the release of IL-10, tumor necrosis factor (TNF)-α, IL-1β, interferon (IFN)-β, transforming growth factor (TGF)-β1 (R&D Systems, Bio-Techne, Novel Châtillon sur Seiche, France) and IL-6 (Clinisciences, Nanterre, France) was quantified using specific immunoassay kits. The sensitivity of the assays was (pg/ml) 15.4 for IL-6, 3.9 for IL-10, 5.5 for TNF-α, 0.125 for IL-1β, 50 for IFN-β and 4.61 for TGF-β1.

### Flow cytometry

PBMCs from healthy donors or Covid-19 patients were resuspended in PBS (Life Technologies) containing 5% FBS and 2mM EDTA (Sigma-Aldrich) for 20 minutes before staining using the following fluorochrome-conjugated antibodies (mouse IgG1): CD3 (UCHT1), CD20 (B9E9), CD14 (RMO52), CD16 (3G8) purchased from Beckman Coulter, Paris, France; HLA-DR (G46-6) and CD163 (GHI/61) from BD Biosciences, Le Pont de Claix, France, and appropriate isotype controls. A minimum of 50,000 events were acquired for each sample using a BD Canto II instrument (BD Biosciences) and data were analyzed with FlowJo software (Tree Star, Ashland, OR).

### Statistical analysis

Statistical analysis was performed with GraphPad Prism (7.0, La Jolla, CA), using the two-way ANOVA test for viral quantification (Ct values) and transcriptional analysis, nonparametric Kruskall-Wallis test for group comparison, nonparametric Mann-Whitney *U* test for cytokine levels, and nonparametric t-test for flow cytometry results with monocyte populations and surface marker expression. Turkey’s and Sidak’s tests were used for post-hoc comparisons. qRT-PCR data for monocytes and macrophages, including principal component analysis (PCA) and hierarchical clustering of gene expression, were analyzed using the ClustVis webtool (23). Differences were considered statistically significant at *P* < 0.05.

## Authorship contributions

A.B, L.G, S.M, A.B.B, M.M and J.V performed the experiments and analyzed the data. S.M, B.D, D.R, B.L.S, P.H, J.V, D.O and J.L.M supervised the work. S.M, J.V and J.L.M wrote the paper. All the authors read and approved the final manuscript.

## Acknowledgments

Dr. Corinne Brunet, Pr. Françoise Dignat-George and Dr. Romaric Lacroix, for assistance with immunophenotyping and sample tracking. Asma Boumaza was supported by the “Fondation Méditerranée Infection". Soraya Mezouar was first supported by the “Fondation pour la Recherche Médicale” postdoctoral fellowship (reference: SPF20151234951) and then by the “Fondation Méditerranée Infection". This work was supported by the French Government under the “Investissements d’avenir” (investments for the future) program managed by the “Agence Nationale de la Recherche” (reference: 10-IAHU-03). The team “Immunity and Cancer” was labeled “Equipe FRM DEQ 201 40329534” (for DO). This work was supported by the IMMUO-COVID project managed by the “Agence Nationale de la Recherche” Flash Covid (reference: IMMUNO-COVID).

