## Supplementary figures and images for "Monocytes and macrophages, targets of SARS-CoV-2: the clue for Covid-19 immunoparalysis"

### Supplemental figure

#### Supplemental figure 1

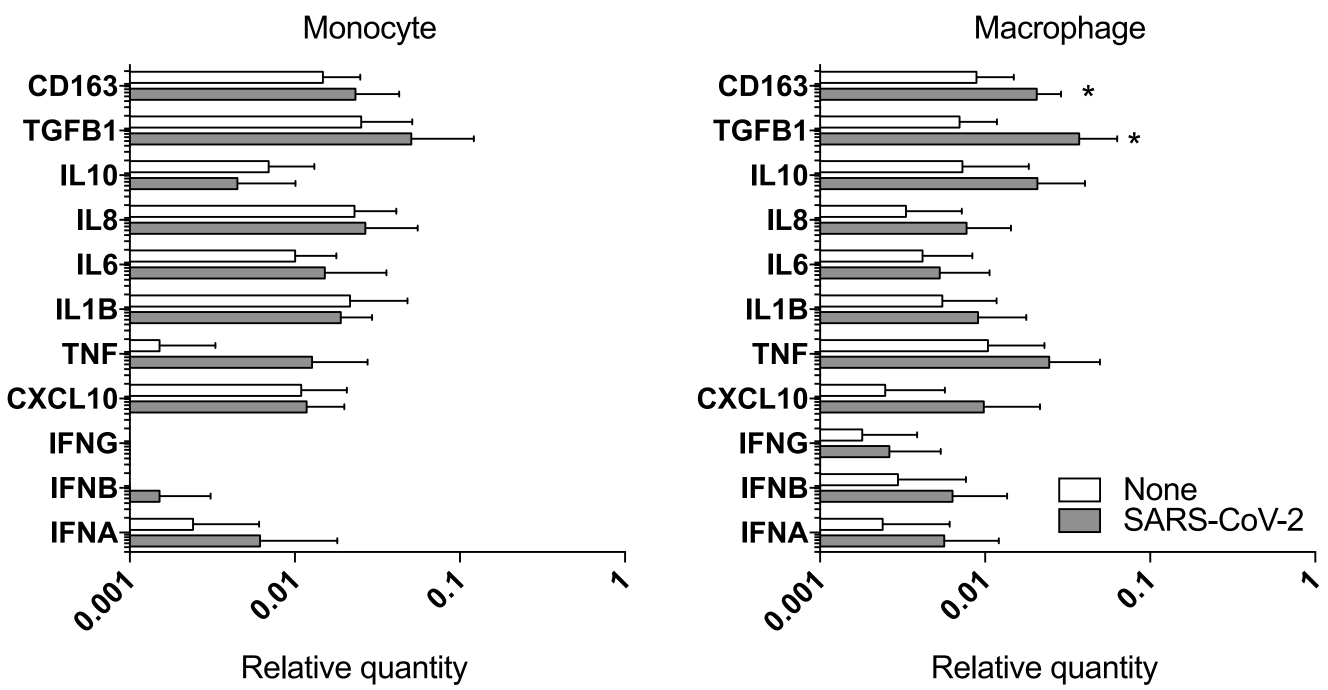

Supplemental figure 2

A.

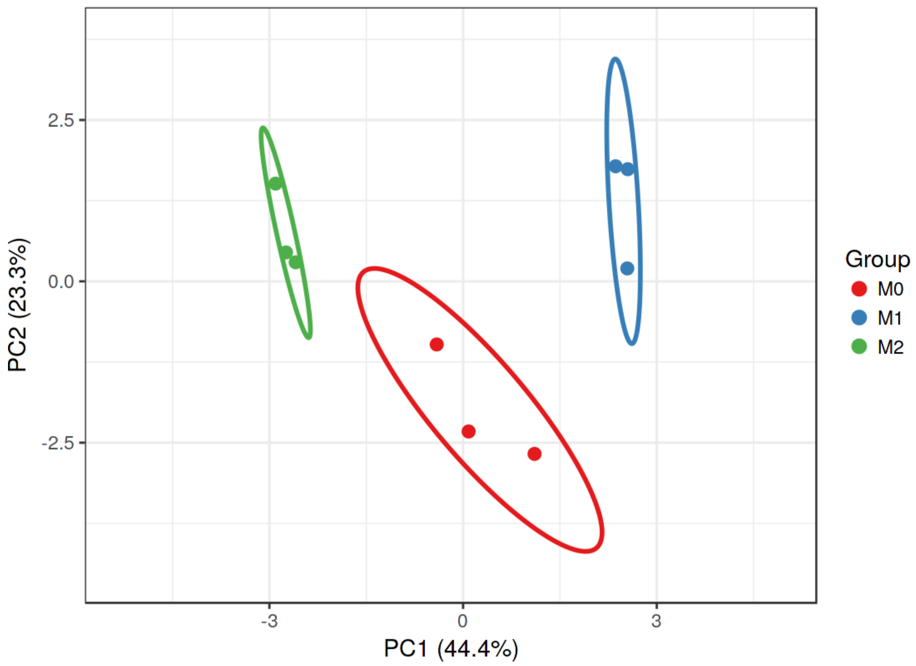


B.

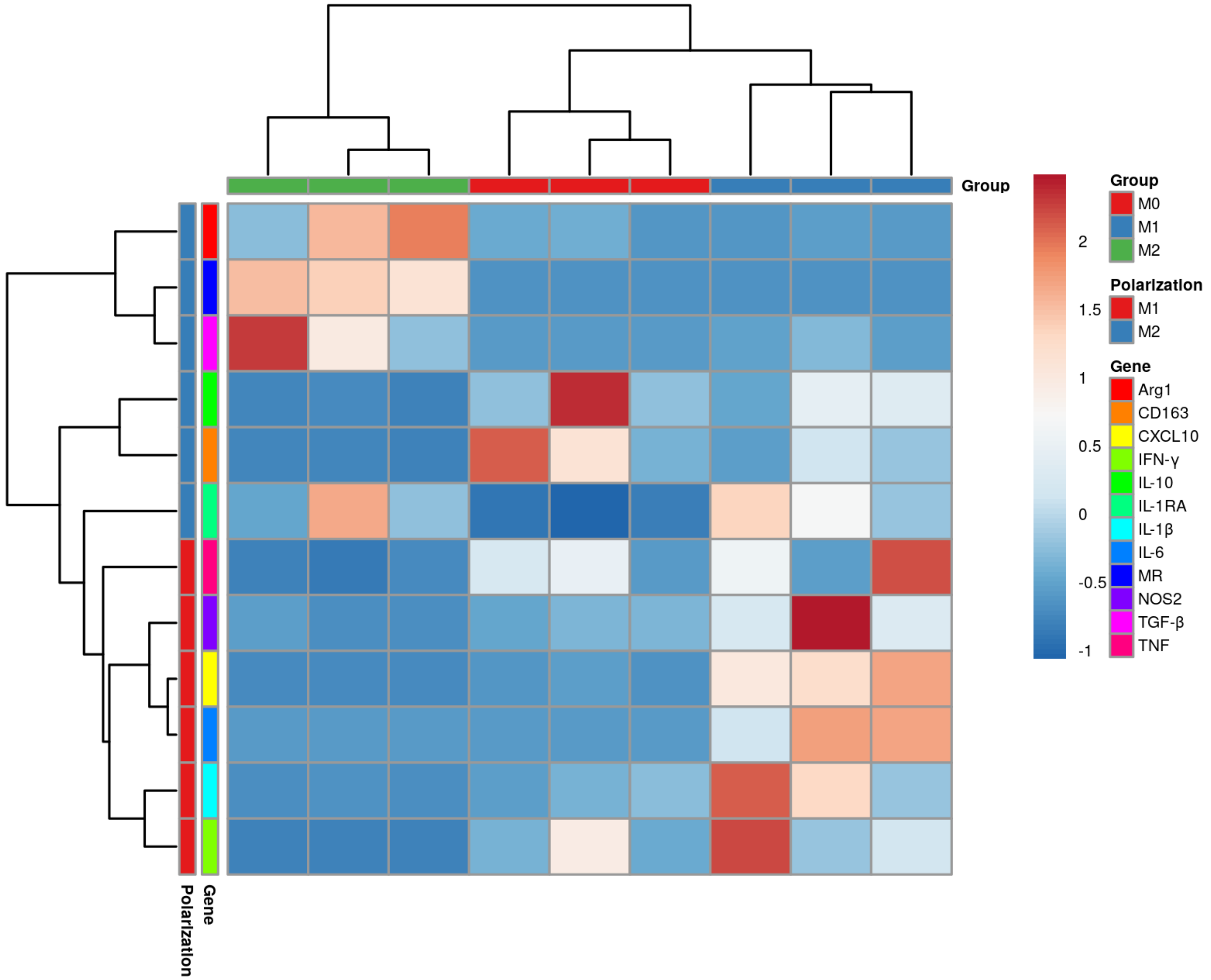

Supplemental figure 3

A.

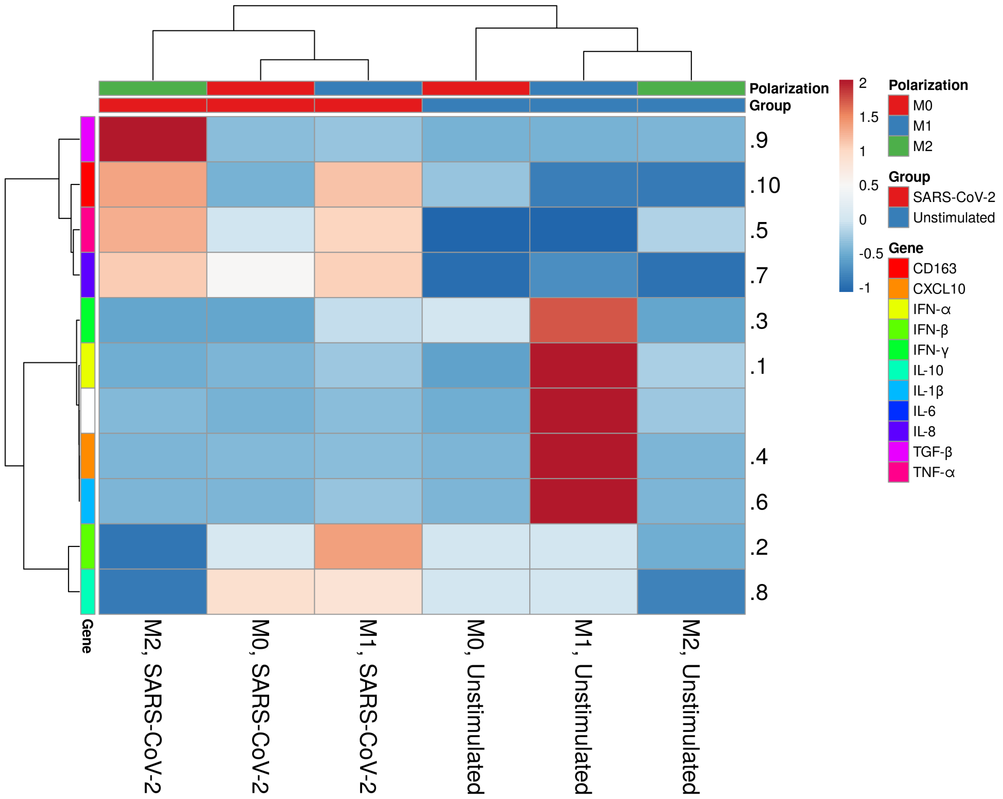


B.

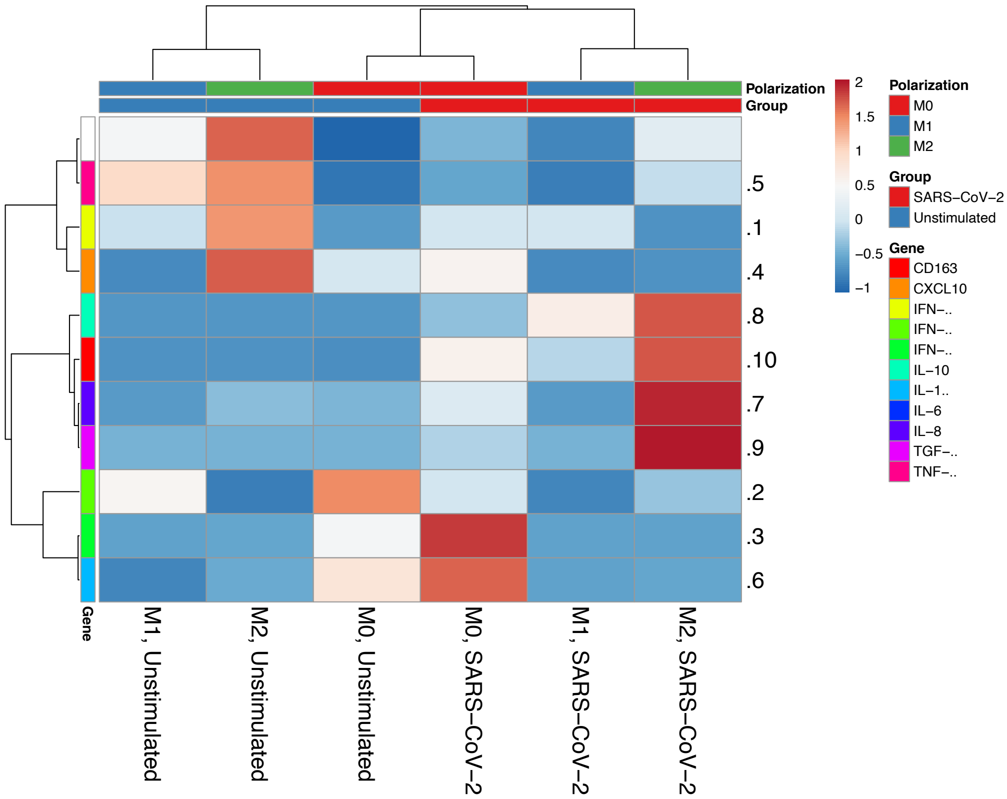
